# Metagenomic reconstruction of nitrogen and carbon cycling pathways in forest soil: Influence by different hardwood tree species

**DOI:** 10.1101/2020.06.23.167700

**Authors:** Charlene N. Kelly, Geoffrey W. Schwaner, Jonathan R. Cumming, Timothy P. Driscoll

**Affiliations:** Division of Forestry and Natural Resources, West Virginia University, Morgantown, West Virginia, USA; National Ecological Observation Network, Aquatics, NEON Domain 07, Oak Ridge, Tennessee, USA; Department of Biology, West Virginia University, Morgantown, West Virginia, USA

## Abstract

The soil microbiome plays an essential role in processing and storage of nitrogen (N) and carbon (C), and is influenced by vegetation above-ground through imparted differences in chemistry, structure, mass of plant litter, root physiology, and dominant mycorrhizal associations. We used shotgun metagenomic sequencing and bioinformatic analysis to quantify the abundance and distribution of gene families involved in soil microbial N and C cycling beneath three deciduous hardwood tree species: ectomycorrhizal (ECM)-associated *Quercus rubra* (red oak), ECM-associated *Castanea dentata* (American chestnut), and arbuscular mycorrhizal (AM)-associated *Prunus serotina* (black cherry). Chestnut exhibited the most distinct soil microbiome of the three species, both functionally and taxonomically, with a general suppression of functional genes in the nitrification, denitrification, and nitrate reduction pathways. These changes were related to low inorganic N availability in chestnut stands as soil was modified by poor, low-N litter quality relative to red oak and black cherry soils.

**IMPORTANCE:** Previous studies have used field biogeochemical process rates, isotopic tracing, and targeted gene abundance measurements to study the influence of tree species on ecosystem N and C dynamics. However, these approaches do not enable a comprehensive systems-level understanding of the relationship between microbial diversity and metabolism of N and C below-ground. We analyzed microbial metagenomes from soils beneath red oak, American chestnut, and black cherry stands and showed that tree species can mediate the abundance of key microbial genes involved in N and (to a lesser extent) C metabolism pathways in soil. Our results highlight the genetic framework underlying tree species’ control over soil microbial communities, and below-ground C and N metabolism, and may enable land managers to select tree species to maximize C and N storage in soils.

## Introduction

Soil communities established among roots, mycorrhizal fungi, and a diverse assemblage of microbiota — the soil microbiome — play pivotal roles in the capture, processing, and storage of environmental **nitrogen** (**N**) and **carbon** (**C**). The size, structure, and function of these communities are affected by changes in the dominant above-ground tree species (1), which influence the pools of available N and C through *a)* alterations in the chemical composition of leaf litter (2–4), *b)* alterations in the composition of root litter and exudates (5, 6), and *c)* the introduction of different mycorrhizal species (7–9). These changes, in turn, can have a significant impact on overall forest productivity (10–12).

One important component of forest productivity is the influence of tree species on below-ground N cycling. Controls on N cycling play a large role in the C budget of ecosystems (12, 13) by influencing C storage both above- and below-ground (14, 15). Different tree species have been associated with divergent rates of ecosystem N cycling and loss (5, 16, 17), and such effects may be mediated by the formation of specific mycorrhizal fungal relationships below-ground (9, 12). For example, soils beneath **arbuscular mycorrhizal** (**AM**)-associated species such as maple (*Acer* spp.), yellow poplar (*Liriodendron tulipifera*), and black cherry (*Prunus serotina*) exhibit increased N mineralization rates and availability of inorganic N relative to soils beneath **ectomycorrhizal** (**ECM**)-associated species such as oak (*Quercus* spp.) and spruce (*Picea* spp.) (18, 19). Changes to N cycling and availability may also arise from differences in soil microbial substrate quality, which has been shown to influence net NO_3_-N production (18).

In some cases, high N availability can lead to a reduced richness and diversity of microbial genes that encode enzymes used to degrade recalcitrant C, leading to an increase in C storage in litter and surface soils (20–22). In other cases, it may lead to greater microbial biomass and byproducts, increasing C storage via protected organo-mineral interactions on clay minerals particularly in subsurface soil (9, 23). These and related processes lead to substantial differences in soil C stocks beneath different tree species; in hardwood systems, these differences can range from minimal to up to 80-ton ha^-1^ when accounting for total C in soil and biomass (14, 24).

Although it is not evident how changes in dominant tree species contribute to long-term N retention and **soil organic matter** (**SOM**) storage, it is clear that changes in dominant above-ground vegetation lead to changes in soil nutrient cycling and C storage through their complex influence on the soil microbial community. Consequently, a systems-level understanding of soil microbiome functional capacity is necessary to fully appreciate the impact of above-ground species shifts that arise in response to ecosystem disruptions such as climate change, harvest, management influences, or species diversity loss via disease (14).

Soil microbial communities contribute to the global N cycle through a collection of interconnected biochemical pathways (25–28), including N mineralization, immobilization, and oxidation-reduction reactions, that interconvert between different species of N (**Fig. 1**). Historically, microbial N cycling has been estimated by measuring biogeochemical process rates of microbial N transformation and/or plant uptake of NO_3_^-^, NH_4_^+^, and organic N (29, 30). More recent studies have used **quantitative polymerase chain reaction** (**qPCR**) to measure the abundance of key functional genes involved in N cycling (28), including *amoA* (nitrification) (31, 32), *narG* (nitrate reduction) (33), *nirK* and *nirS* (nitrite reduction) (34), *norB* (NO reduction) (35), *nosZ* (N_2_O reduction) (36), and *nifH* (N_2_ fixation) (37). A growing body of evidence supports functional gene abundance as a useful index for predicting biogeochemical rates, particularly when applied to N process rates (33, 38–40).

**Fig. 1.**
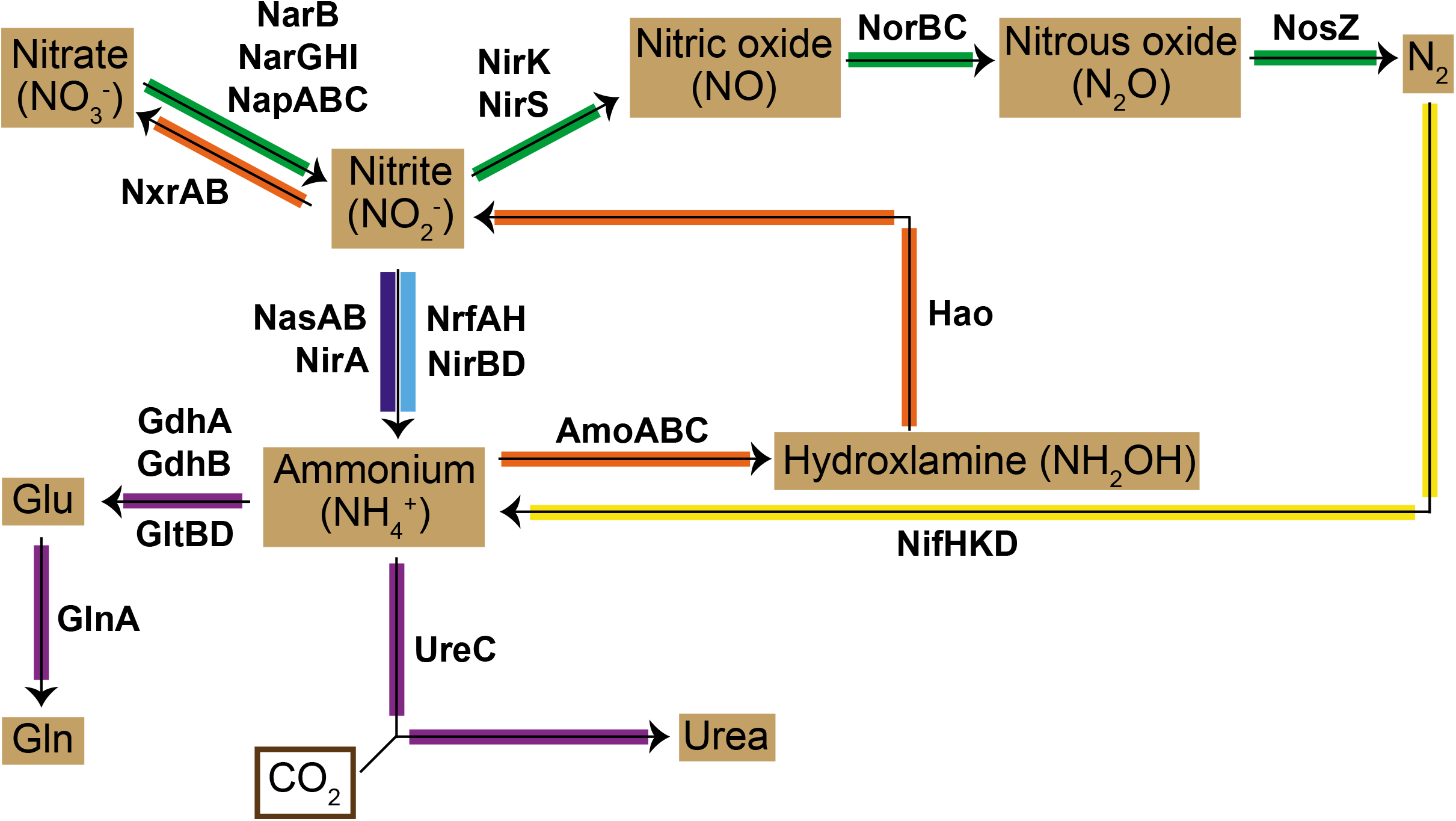
Nitrogen metabolism pathway illustrating gene families responsible for transformation between N species.

In addition to N cycling, microbial communities also play an important role in soil C dynamics and storage (41) through their varied roles as scavengers, detritivores, plant symbionts, and pathogens. Differences in the functional roles of microbes determine nutrient availability, organic matter turnover, and ultimate C retention in soil (42–44) through differences in production of various extracellular enzymes used in C and nutrient scavenging and acquisition. There are few studies that have investigated the link between soil microbial genomic structure and the C cycle, although one such study (44) reported a strong relationship between functional genes or exoenzymes (including acetylglucosaminidase, amylase, and xylanase) and corresponding enzyme activities related to C degradation (N-acetyl-β-D-glucosaminide, a-D-glucopyranoside, and β-D-xylopyranoside, respectively).

The characterization of functional genes in complex ecosystems traditionally has relied on primers to amplify and quantify selected gene targets. These approaches have yielded important insights; however, they do not provide a comprehensive view of microbial genetic capacity, nor are such studies able to assess the contribution of novel or uncultured species in ecosystem functioning. Recent advances in high-throughput DNA-based methods (e.g., metagenomics) have enabled the unbiased interrogation of entire soil microbial communities and have been applied to the functional characterization of soil microbiota in complex ecosystems, particularly in response to changing conditions such as elevated CO_2_ (28). However, we know of no studies that have leveraged metagenomics to assess the effect of above-ground tree species on the functional profiles of below-ground microbial communities.

In the present study, we used shotgun metagenomic sequencing to quantify the abundance of gene families involved in soil microbial N and C cycling beneath replicated plots of three deciduous hardwood tree species: ECM-associated *Quercus rubra* (red oak), ECM-associated *Castanea dentata* (American chestnut), and AM-associated *Prunus serotina* (black cherry). We hypothesized that gene abundance related to nitrification and denitrification in particular would be elevated in soil beneath cherry, due to its AM association and high N mineralization, and that these same genes would occur in lesser abundance in soil beneath chestnut, due to its ECM association and low N mineralization.

## Methods

### Study design

This study was conducted at Purdue University’s Martell Research Forest in West Lafayette, Indiana, USA (40° 26’ 42” N, 87° 01’ 47” W). This 2.4-hectare (ha) plantation of pure, non-hybrid northern red oak, American chestnut, and black cherry was planted in 2007 to study chestnut growth dynamics (45). Prior to 2007, plots were maintained in corn production. The plantation is comprised of seven species compositions: three pure stands (one for each species), three mixed stands of all species pair combinations, and a single mixed stand containing all three species. The present study was restricted to soil from pure stands of the three species, with tree spacing regimes of 1×1 meter (10,000 stems ha^-1^, 5×5 meter plot size) and 2×2 meter (2,500 stems ha^-1^, 10×10 meter plot size) spacing, resulting in eight plots comprising an experimental block. and replicated three times at the site (3 species, 2 densities, 3 replications; N = 18 plots). At the time of soil collection, trees were 10 years old. Mean tree **diameter at breast height** (**DBH**) was 7.10, 8.23, and 8.36 centimeters, and mean tree height was, and 6.53, 6.49, and 4.46 meters for oak, chestnut, and cherry, respectively, across all plots. Plots were typically under closed canopy and all soil samples were collected beneath live, healthy trees within each plot.

Sampled soils are the Rockfield series: mildly acidic to nearly neutral, mainly silt loams with a clay content of approximately 20–32% (Soil Survey Staff), moderately productive, deep, and formed from silty outwash and loamy till. Soil profiles were similar at all three blocks, showing a 2–5 cm Ap-horizon (agriculturally disturbed), weak Bt-horizon development, and no significant surface organic O-horizon development. There were no measurable differences in current total soil C content, N content, or soil pH between species or spacing treatments, as analyzed by horizon (Ap, Bt1, Bt2) in 2017. Mean soil C across the plots was 5866 kg ha^-1^ and soil pH in the Ap-horizon was 5.76. From 1981–2010, mean annual temperature was 10.4° C and mean annual precipitation was 970 mm (National Climatic Data Center 2018).

### DNA extraction

Soil samples were collected in June 2017 with each sample composited from four soil cores (0–15 cm) from each plot. Soils were transported on ice to the laboratory, immediately sieved to 2 mm, and stored at –80° C. DNA was extracted using the DNeasy PowerSoil kit (Qiagen) with the following modifications: 0.5 g of soil was suspended in 500 μl of DNeasy PowerSoil lysis buffer, 250 μl of 0.2 mm stainless steel beads, and five 3.2-mm stainless steel beads, then homogenized for 5 minutes using a Bullet Blender tissue homogenizer (NextAdvance), centrifuged at 12,000 rpm for 8 minutes, and the supernatant used for DNA extraction. DNA was quantified by fluorimetry (Qubit 3.0, Invitrogen) and assessed for quality using a Nanodrop spectrophotometer (Thermo Scientific).

### Metagenome sequencing and bioinformatic analysis

Approximately 500 ng of DNA from each sample (n=18) was submitted to the Department of Energy’s **Joint Genome Institute** (**JGI**) for metagenome sequencing, contig assembly, annotation, and gene copy number determination as described previously (46). Processed data were deposited in the **Integrated Microbial Genomes/Microbiomes** (**IMG/M**) database (**Table S1**). Functional gene abundance values for each metagenome were calculated as copies per 10^6^ total protein **coding sequences** (**CDS**) to allow for comparison across metagenomes of different sizes, and averaged across replicate samples. Gene families involved in N and C cycling were identified using a combination of **Kyoto Encyclopedia of Genes and Genomes** (**KEGG**) and **Enzyme Classification** (**EC**) identifiers. Statistical significance of differences in gene abundance between metagenomes was assessed with one-way **analysis of variance** (**ANOVA**) and Tukey multiple comparisons of means, following Shapiro-Wilk test for normality, using SAS JMP version 11.0.0 (SAS Institute, Cary, NC). Microbial diversity was assessed using total CDS assigned across 95 taxonomic classifications in IMG/M and normalized to 10^6^ total metagenome CDS as described for functional gene analysis. Overall differences between microbial communities under different tree species were analyzed using **principal components analysis** (**PCA**) at the level of taxonomic superkingdom (archaea, bacteria, eukaryota, and virus) using the prcomp method in R v3.5.1.

## Results

Here we present our results in the context of microbial gene families that comprise seven N cycle pathways (28): nitrification, denitrification, **assimilatory nitrate reduction** (**ANRA**), **dissimilatory nitrate reduction** (**DNRA**), **anerobic ammonium oxidation (anammox)**, N_2_ fixation, and organic N metabolism (**Fig. 1**). We additionally report abundance data for nitrate and nitrite transport genes, C metabolism genes, and twelve genes involved in stress responses. The KEGG and EC numbers used to identify these gene families are shown in **Table S6**.

### Nitrogen cycle pathways

Overall, soils beneath chestnut showed significantly reduced denitrification and DNRA gene abundances compared to both oak and cherry. Mean abundance of nitrification genes was very low (<1 copy/10^6^ CDS) regardless of tree species, and no genes involved in anammox were identified in any sample. Genes involved in N_2_ fixation were also low in all soils regardless of tree species.

#### Nitrification

The conversion of NH_4_^+^ to NO_3_^-^ proceeds via three oxidation steps catalyzed by the products of *amoABC* (ammonia monooxygenase), *hao* (hydroxylamine dehydrogenase), and *nxrAB* (nitrite oxidoreductase) gene families. Abundances of *amoABC* and *hao* gene families were consistently low (<5 copies/10^6^ CDS) to undetectable across all soils regardless of dominant tree species (**Fig. 2 [orange], Table S2A**). We were unable to determine the abundance of *nxrA* or *nxrB* directly, as IMG/M does not distinguish between *nxrAB* and *narGH;* consequently, we reference abundance values for *narG* and *narH* instead.

**Fig. 2.**
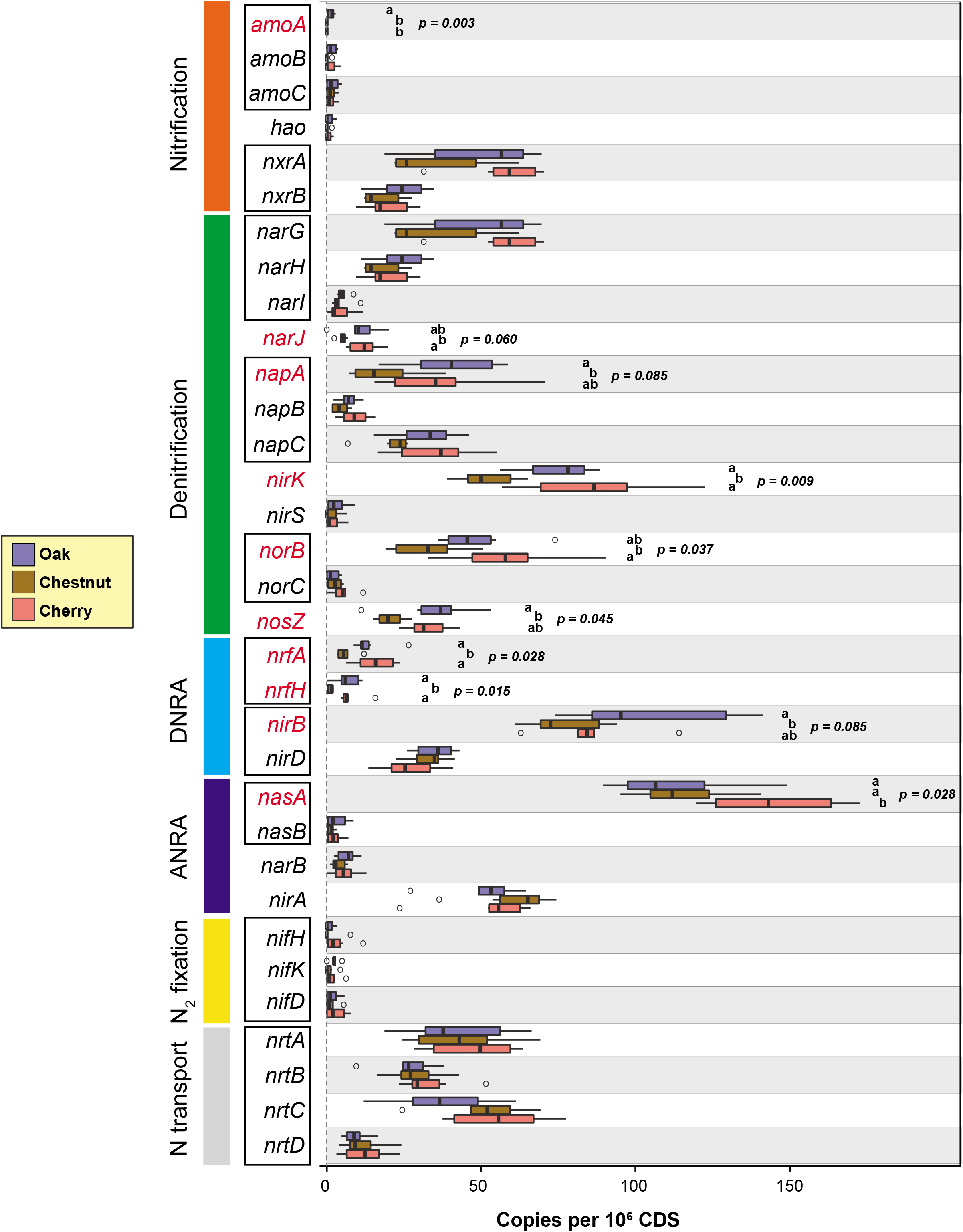
Mean abundance values (n=6) of gene families involved in nitrification (orange), denitrification (green), DNRA (light blue), ANRA (dark blue), N_2_ fixation (yellow), and N transport (gray). The total number of genes in each sample was normalized by the number of protein-coding genes in the genome and scaled to 10^6^. Red text denotes genes with a significant difference in abundance by tree species (p<0.1 using ANOVA and Tukey’s; see Methods). Blue: soils beneath oak; orange: soils beneath chestnut; red: soils beneath cherry. See **Table S6** for database accession identifiers for each gene.

#### Denitrification

The reduction of NO_3_^-^ to N_2_ gas proceeds via four steps and produces several nitrogenous compounds with notable roles as air-polluting gases, including N_2_O and NO. Twelve microbial gene families involved in denitrification were included in our analysis, all of which were detectable in most samples (**Table S2B**). Overall, genes involved in denitrification were substantially reduced in soils beneath chestnut compared to both oak and cherry, even after normalization to total CDS (**Fig. 2 [green]**). Gene families involved in the reduction of NO_2_^-^ to NO (*nirS, nirK*) and NO to N_2_O (*norBC*) were significantly higher in soils beneath cherry compared to either ECM-associated species, although the abundance of *nirS* was very low to undetectable in many samples regardless of species. *nosZ*, the principal gene involved in reduction of N_2_O to N_2_, was elevated in soils beneath oak compared to chestnut.

#### Nitrate reduction

The reduction of NO_3_^-^ to NH_4_^+^ ultimately leads to the incorporation of N into microbial biomass. DNRA is an anaerobic process in which NO_3_^-^ is used as an electron acceptor to oxidize and release energy from organic C. It is mediated by nitrate reductases (*narGHI, napABC*) that form NO_2_^-^ and nitrite reductases (*nrfAH, nirBD*) that convert NO_2_^-^ to NH_4_^+^. In all samples *nirBD* complex genes were consistently 2-to 10-fold more abundant than *nrfAH*, regardless of species (**Fig. 2 [light blue], Table S2C**). In contrast, *nrfAH* genes were significantly reduced in soils beneath chestnut compared to oak and cherry. Unlike DNRA, ANRA is an energetically costly process and involves different families of nitrate (*nasAB, narB*) and nitrite (*nirA*) reductases. The nitrate reductase catalytic subunit (*nasA*) was predominant in all samples, and significantly enriched in soils beneath AM-associated cherry trees. In contrast, the nitrite reductase *nirA* was slightly more abundant in chestnut-associated soils (**Fig. 2 [dark blue]**).

#### Annamox

Anaerobic ammonium oxidation is the process by which NH_4_^+^ and NO_2_^-^ are metabolically combined to form N_2_ gas. It is mediated by the gene families *hzsA* (hydrazine synthesis) and *hzo* (hydrazine oxidoreductase). Neither *hzsA* nor *hzo* were detected in the metagenomes of any of our soils (**Table S2D**), indicating no capacity for hydrazine synthesis and anaerobic oxidation of ammonia.

#### Nitrogen fixation

The reduction of N_2_ gas to biologically available NH_4_^+^ is carried out by the nitrogenase complex encoded in the *nifDKH* operon. Abundance of these genes was measurable but consistently low (<5 copies per 10^6^ CDS) across all samples, regardless of tree species (**Fig. 2 [yellow], Table S2D**).

#### Nitrogen transport

The *nrtABCD* gene cluster encodes an **ATP-binding cassette** (**ABC**)-type transporter capable of importing NO_3_^-^ or NO_2_^-^ from the extracellular environment. Genes in this cluster were detected in all samples (**Fig. 2 [gray], Table S2D**) and were marginally more abundant in soils beneath AM-associated cherry trees.

#### Organic N metabolism

Incorporation into microbial biomass is another important potential fate of soil N. Here we assayed several different pathways for organic N metabolism, including the conversion of NH_4_^+^ to glutamate (*gltBD, gdhA, gdhB*), glutamine (*glnA*), and urea (*ureC*). We observed no significant difference in the abundance of these genes between different tree species (**Fig. 3, Table S3**). The gene responsible for dissimilatory glutamate formation (*gdhB*) was markedly lower than other families across all metagenomes. Additionally, *nao* (nitroalkane oxidase) and *nmo* (nitronate monooxygenase) were absent from all samples, indicating that the degradation of nitroalkane compounds was not a significant source of N in these soils. The regulatory gene *glnR*, which represses glutamine synthesis from glutamate by *glnA*, was highly abundant in all soils.

**Fig. 3.**
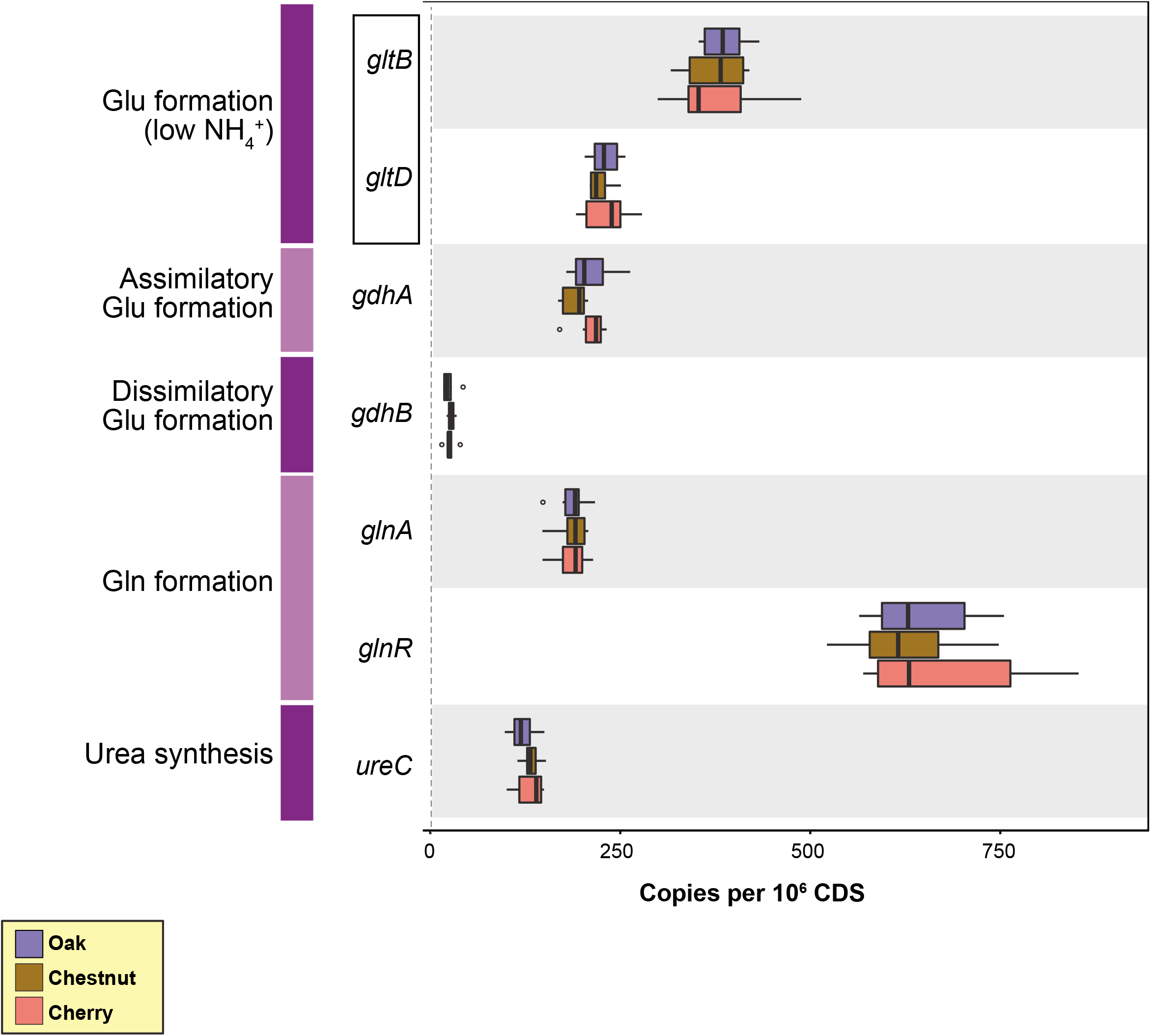
Mean abundance values (n=6) of gene families involved in organic N decomposition. Data normalization and color scheme according to **Fig. 2.**

### Carbon metabolism genes

We analyzed 15 gene families encoding enzymes for C degradation, grouped by the type of compound they metabolize: starch, hemicellulose, cellulose, aromatics, chitin, and lignin (**Fig. 4, Table S4**). The most abundant C-metabolizing gene family across all soils was β-glucosidase (hemicellulose-degrading), roughly twice as prevalent than the next most abundant family, α-amylase (starch-degrading). Although high, the abundance of β-glucosidase was somewhat reduced in soils beneath cherry compared to both ECM species. Also reduced in soils beneath cherry was vanillate monooxygenase, an exoenzyme that degrades moderately recalcitrant aromatic compounds. In contrast, chitinase was slightly more abundant in cherry-associated soils compared to oak, with chestnut-associated soils containing intermediate levels. The cellulose degrading 1,4-β-cellobiosidase was also reduced beneath oak relative to both chestnut and cherry.

**Fig. 4.**
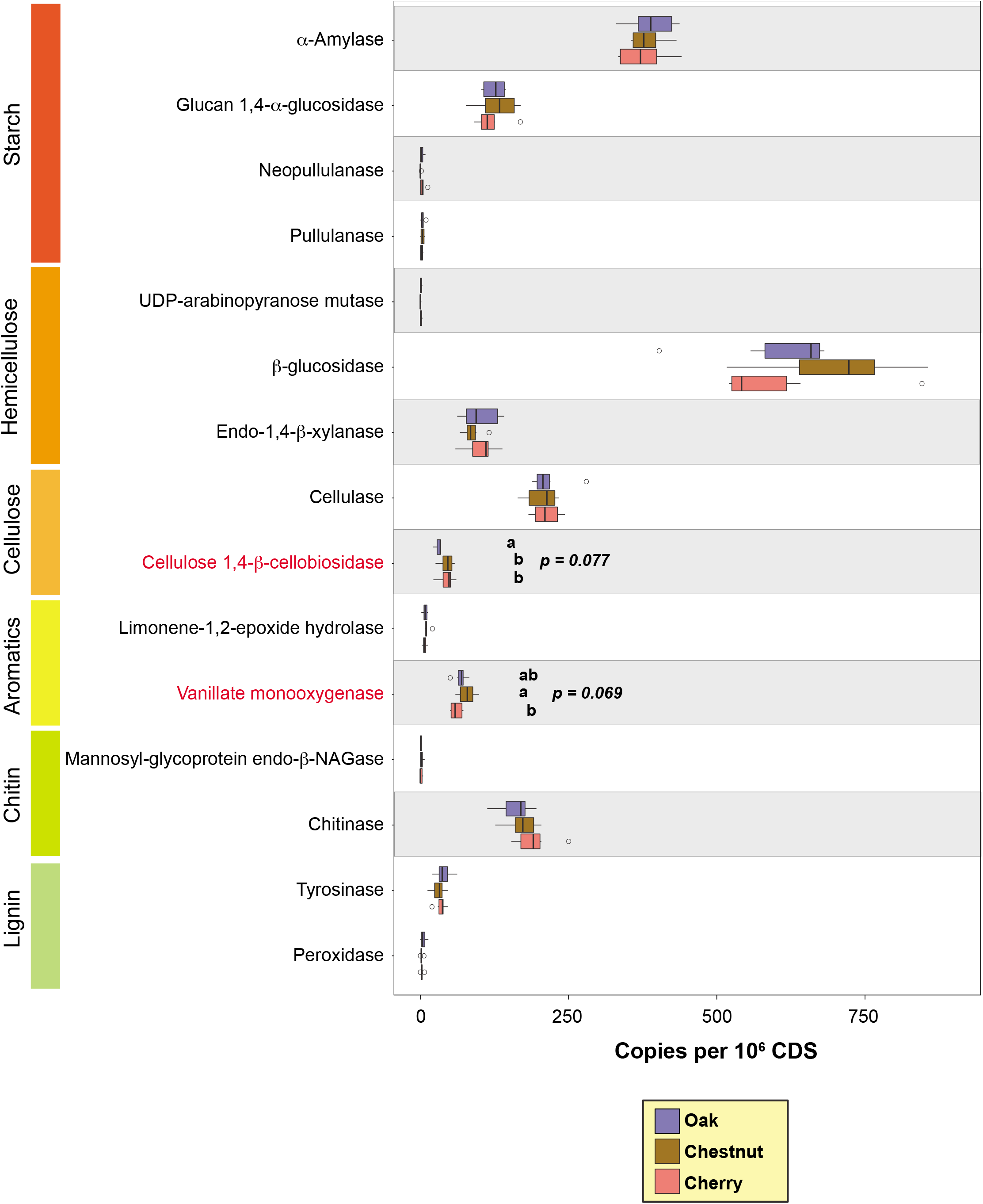
Mean abundance values (n=6) of gene families involved in carbon metabolism. Data normalization and color scheme according to **Fig. 2.**

### Stress response genes

We assayed a total of 12 gene families encoding enzymes related to general stress responses and found soils beneath chestnut trees had significantly higher numbers of choline dehydrogenase and catalase genes, and significantly reduced abundance of gentisate 1,2-dioxygenase and superoxide dismutase genes compared to cherry (**Fig. 5, Table S5**). No other differences between tree species were observed. Glycosylases were the most abundant stress response family across all soils.

**Fig. 5.**
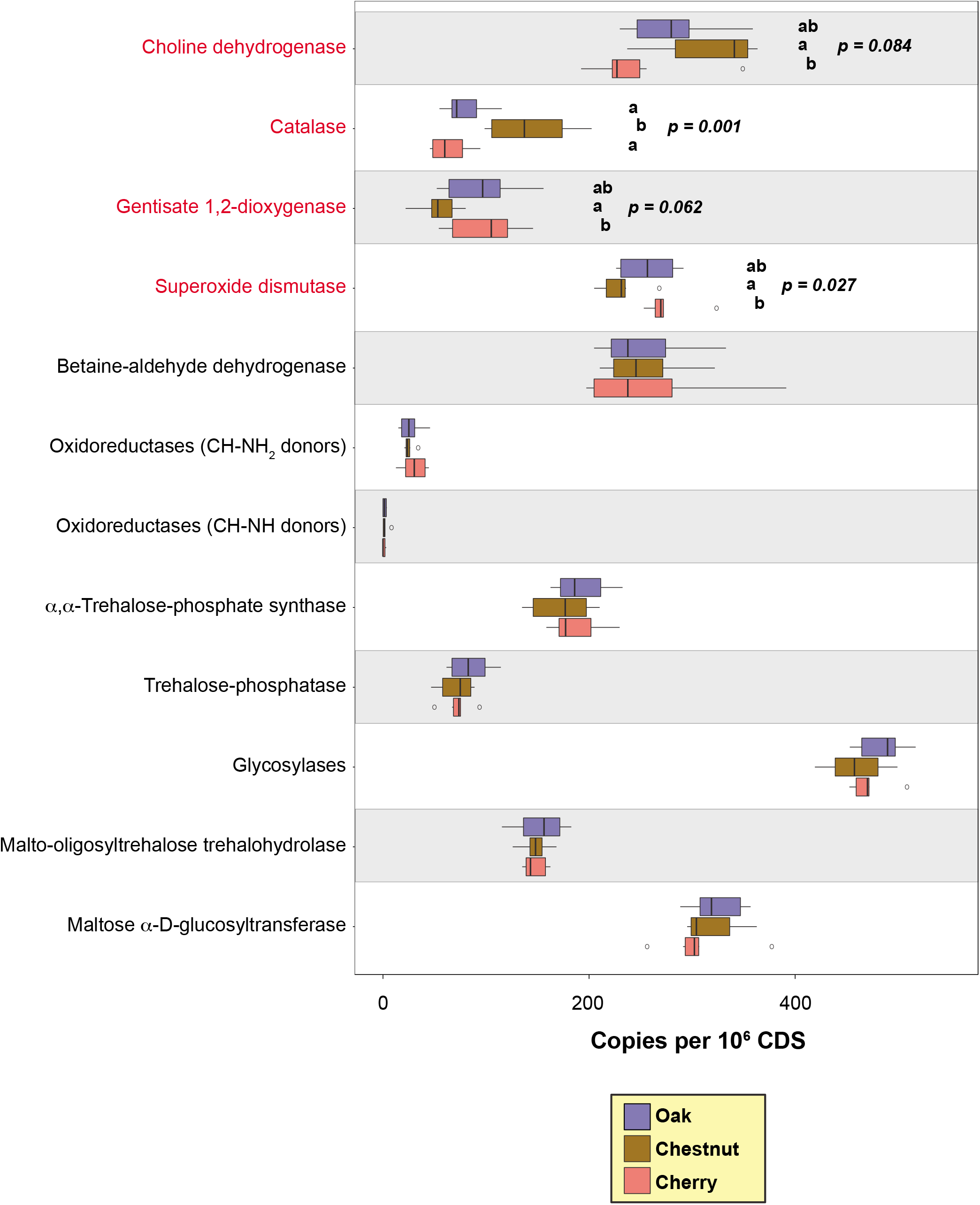
Mean abundance values (n=6) of gene families involved in stress response. Data normalization and color scheme according to **Fig. 2.**

### Taxonomic distribution

Analysis of all microbial taxonomic groups by PCA revealed distinct separation between cherry-associated and chestnut-associated soil metagenomes (**Fig. 6A**); oak-associated soils were intermediate but slightly more aligned with cherry. A similar pattern was seen when taxa included only bacterial (**Fig. 6B**) and only eukaryotic groups (**Fig. 6C**). Viral (**Fig. 6D**) and archaeal (**Fig. 6E**) subsets were more homogeneous, possibly a consequence of their smaller group sizes (n=5 and n=8, respectively).

**Fig. 6.**
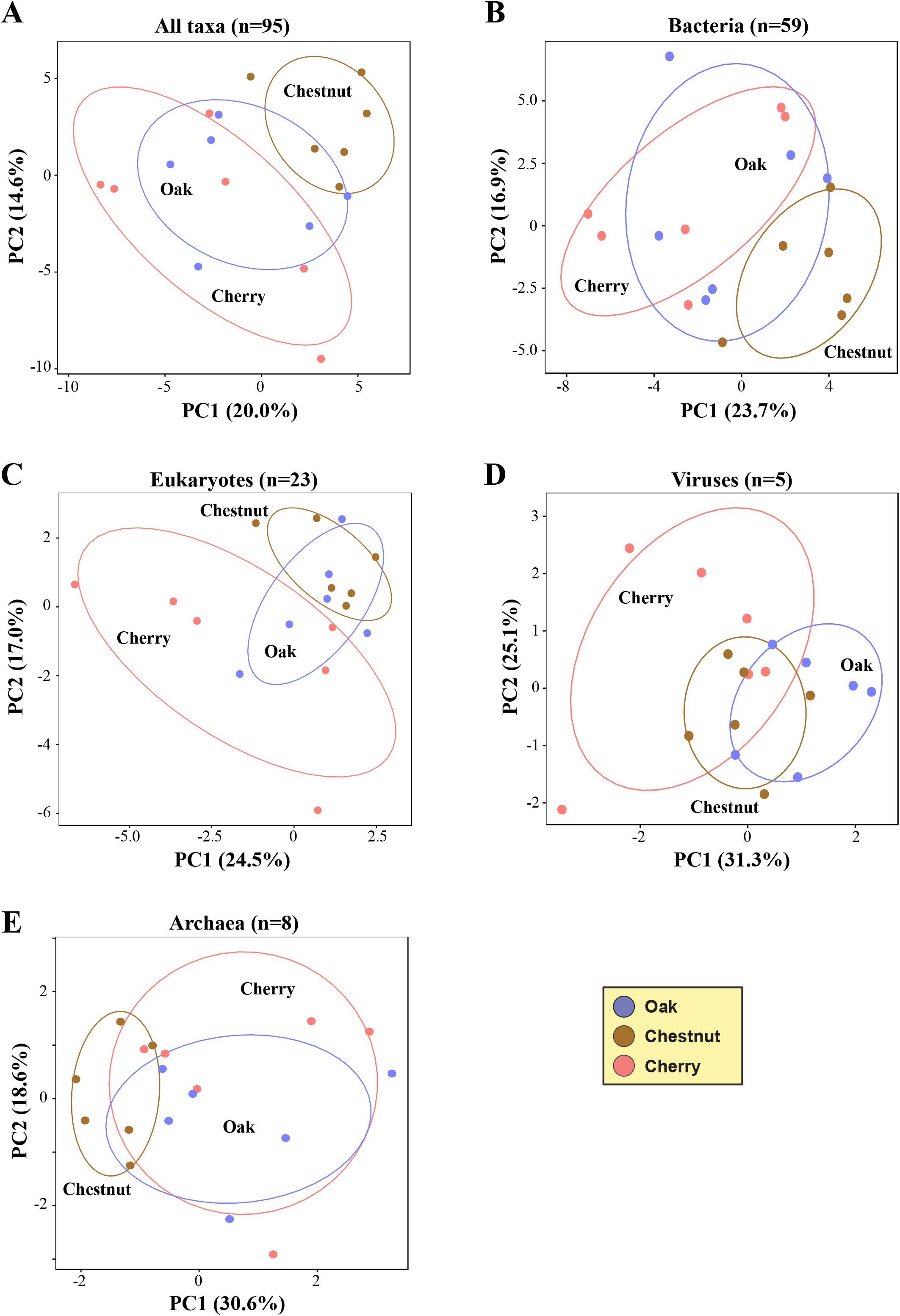
Principal components analysis of taxonomic groups occurring within all metagenomes (n=18) and clustered by tree species. Blue: soils beneath oak; orange: soils beneath chestnut; red: soils beneath cherry. **A**. All taxa combined (n=95). **B**. Bacterial taxa (n=56). **C**. Eukaryotic taxa (n=23). **D**. Viral taxa (n=5). **E**. Archaeal taxa (n=8).

**Table 1.**
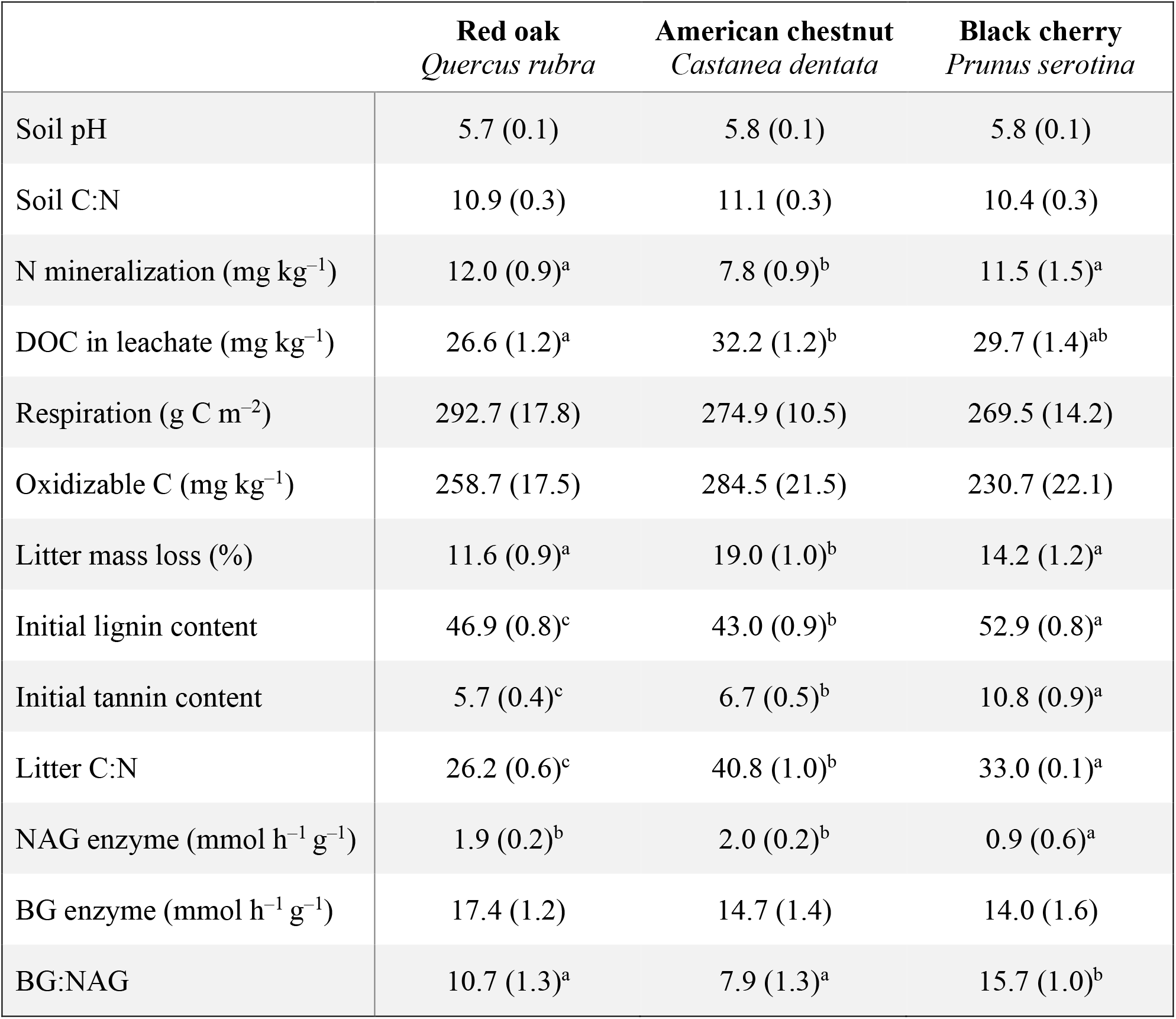
Parameters and process rates from a one-year incubation of litter and soil associated with three tree species. Adapted from Schwaner & Kelly (2019). Superscript letters indicated significant difference (p<0.05) by ANOVA and Tukey’s HSD; see Methods for details.

## Discussion

Changes in dominant tree species are expected to occur in many locations due to climate change (47) and deliberate species introductions in plantation and restoration efforts (48). These changes are likely to alter soil microbial communities and the associated C and N dynamics below-ground; however, the extent and mechanism by which tree species exert influence on particular functional gene families involved in N and C metabolism is still poorly understood. In this study, we apply a combination metagenome sequencing and bioinformatic analysis to assess, for the first time, the impact of tree species differences on functional gene abundances in the soil microbiome in a comprehensive and unbiased manner. Overall, we show that tree species can mediate the abundance of multiple key functional genes that encode for enzyme production in N and C metabolic pathways, and soils beneath chestnut exhibited the most distinct soil microbiomes, both functionally and taxonomically. The results of our study may facilitate the selection of tree species for land management purposes of N management and C storage.

### Nitrogen cycle pathways

We identified significant differences in functional gene abundance as a result of tree species within the nitrification, denitrification, DNRA, and ANRA pathways (**Fig. 2**). Overall, gene abundance exhibited the greatest differences between soils beneath chestnut and cherry, while the gene abundance in soils beneath oak was generally intermediate between cherry and chestnut. This is supported by the significantly lower gene abundance values of *amoA, narJ, nirK, norB, nosZ, narJ, napA, nirB, nrfA, nrfH*, and *nasA* beneath chestnut relative to cherry. This data supports the stated hypothesis that abundance of functional genes related to nitrification and denitrification, in particular, would be elevated in soil beneath cherry. The soils beneath cherry exhibited the greatest N mineralization rates and NO_3_^-^ in leachate, indicating a rapid N cycle and high inorganic N availability (49). As other studies support, we relate the high inorganic N availability in cherry soil to its AM fungal association and relatively low C:N litter, as well as low NAG enzyme activity, indicating that N is not limiting microbial processes in soils beneath cherry.

Abundances of all genes within the nitrification pathway (*amoABC, hao*) were low, regardless of tree species, and *amoA* was not detected in either chestnut or cherry soils (**Fig. 2**). This is contrary to our hypothesis that cherry soil would have greater abundance of functional genes related to nitrification, relative to chestnut or cherry, which are both ECM-associated tree species. That no *amoA* was detected is surprising, as others have shown high numbers of gene copies from forest soils (up to 1.2 x 10^9^ copies per g dry soil of AOA amoA) (50) and others have shown a strong relationship between NO_3_^-^ production and *amoA* gene abundance (38). NO_3_^-^ production was indeed detected in our prior incubation study for soil and litter from each tree species. However, low *amoA* gene abundance from grassland soil has also been reported (28), with a much greater abundance of *nxrA* and *nxrB* gene copies within the nitrification pathway. Our analysis did not include *nxr* genes, as it was not recognized in the IMG/M gene or function processing, but considering other nitrite oxidoreductase genes (*narGH* genes shown in the denitrification pathway here) suggest no difference in nitrite oxidoreductase gene abundance as a function of tree species.

Within the denitrification pathway, chestnut soils had the lowest gene abundance for *narJ, nirK, norB*, and *nosZ* relative to cherry or oak. Low denitrification-related gene numbers in chestnut soil was expected and aligns with the low inorganic N availability in chestnut soils, as denitrification rates are low in N-limited soil (51). N limitation in chestnut soil is likely a function of the high C:N ratio of the leaf litter and evidenced by the higher NAG enzyme activity. *nirK* and *norB* gene abundance was greatest in cherry soil, while *narJ* and *nosZ* were similar in soils beneath cherry and oak. High *nirK* and *norB* in cherry soil, which encode for the reduction of nitrite and nitric oxide, respectively, supports our hypothesis that cherry, with relatively high inorganic N availability, would stimulate denitrification capacity. Additionally, an analysis of the *nirK:nosZ* ratio, which has been used to estimate endpoint N_2_O production from soil (N_2_O:N_2_ ratio) (33, 40), suggests similar ratios across tree species and no difference in the capacity of these particular tree species to influence N_2_O emission relative to N_2_ gas through denitrification processes. Abundance of *nirK* was also much higher (10-fold or more) than *nirS* in all soils. *nirS:nirK* ratio may be related to the N_2_O sink capacity of soil, with a higher N_2_O consumption with increasing *nirS:nirK* ratio, related to the observation that *nosZ* occurs less frequently in genomes of strains with *nirK* relative to those with *nirS* (52).

Within the DNRA pathway, chestnut soil again exhibited the lowest abundance of several functional genes, including *narJ, napA, nirB, nrfA*, and *nrfH* relative to cherry or oak. The DNRA process is thought to play a role in N retention in forest soils by reducing NO_3_^-^ to NH_4_^+^ (53). The amount of inorganic N was consistently low in soil leachate from our incubation study from chestnut soil (49) and the relatively low abundance of DNRA-related genes in chestnut soil suggests low initial inorganic N availability through low C:N litter inputs, rather than N retention through DNRA processes. Relative abundance of NRA genes (greatest abundance of *nirB* nitrite reductase in DNRA and *nasA* nitrate reductase in ANRA, regardless of tree species) aligns with those from a metagenomic assessment of a Minnesota, USA grassland soil (28), but do not reflect those from a GeoChip microarray assessment of a Tennessee, USA forest soil (54), which exhibited greatest relative abundance of *narG* nitrate reductase in the DNRA pathway. Differences in relative abundance of these genes by location may be a function of the two methodologies (metagenomic and microarray) (55).

Within ANRA, *nasA* was most abundant for all tree species, and was significantly more abundant in cherry relative to either chestnut or oak. High *nasA* abundance in soils beneath cherry may be related to greater amount of NO_3_^-^ available to enter the process of assimilatory reduction to NH_4_^+^ for microbial biosynthesis (56) and ANRA is generally low when NH3 is present (57). Greater abundance of *nasA* beneath cherry shown here fits within the mycorrhizal framework of nutrient cycling as cherry, an AM fungal-associated species, should engender greater microbial assimilation of nutrients and microbial biomass relative to ECM-associated species (12, 23).

As expected for these soils influenced by forest tree species, genes related to organic N metabolism were abundant (up to 677 normalized gene copies for *glnR)*, but surprisingly did not vary by tree species. We expected variation in gene abundance related organic N metabolism, particularly *glnR*, as *glnR* has been referred to as the central regulator of N metabolism and mediates the integration of C and N metabolism through uptake and utilization of non-phosphotransferase-system C compounds (58). There is evidence that *glnR* is inhibited in systems with N limitation (59) and is activated when N availability is high (60). However, others suggest that the rate-limiting step of organic matter decomposition is not the mineralization of amino acid to NH_4_^+^, but rather the previous step of depolymerization of proteins to oligopeptides (61–63). In the present soils, even given the relative differences in inorganic N availability, no difference in organic N metabolism gene abundance was detected and differences exhibited in N metabolism support that N metabolism is mediated by protein depolymerization.

For N_2_ fixation, gene abundance for *nifH* was low and uniform between tree species. This is consistent with the low *nifH* abundance reported elsewhere (28) from grassland soils, although they also showed a significant increase of *nifH* with elevated CO_2_, indicating *nifH* abundance can increase with N limitation. Others have also shown *nifH* gene pools to be sensitive to soil properties and plant cover, including total C, N, soil texture, and controls on inorganic N availability (64). That *nifH* abundance was low and uniform between tree species in the present study may be a function of the monoculture plantings and limited understory plant species present that are capable of forming N fixation symbioses and that the free-living N_2_ fixing species (e.g., cyanobacteria, Azotobacter, Clostridium) were rare.

### C metabolism genes, C degradation, and stress response

Between tree species, we document statistically significant differences of abundance of genes responsible for enzyme production to degrade starch (neopullanase), hemicellulose (β-glucosidase), and aromatic (vanillate) compounds. No differences occurred in cellulose, chitin, or lignin-degrading enzymes between tree species. It is surprising that the C-degradation pathways were so similar between tree species given the differences in mycorrhizal associations and litter quality (tannin, lignin, C:N ratio) among tree species. Our hypothesis that the two ECM-associated tree species (chestnut, oak) would support a soil microbial community with high abundance of genes that encode for degradation of aromatic and lignin compounds, and the AM species (cherry) would support a high abundance of genes that encode for degradation of starch and hemicellulose compounds, was not strongly supported. While results show that chestnut soil had the greatest abundance for β-glucosidase and vanillate and the lowest abundance of neopullanase, difference in abundance of these were slight relative to soils beneath oak and cherry. Linking enzyme production to microbial community function remains challenging due the influence of “cheater” microbes that can utilize the byproducts of enzymatic breakdown, but do not synthesize the enzymes themselves (41), as well as the potential inhibition of enzyme activity by tannin compounds (65). However, the minimal differences in genes responsible for enzyme production for C degradation exhibited here are reflected in the soil incubation results showing similar total soil C, oxidizable C, C-degrading enzyme activity, and CO_2_ respiration between tree species (49).

Four genes related to stress response exhibited variation by tree species, including choline dehydrogenase, catalase, gentisate dioxygenase, and superoxide dismutase, each of which are considered oxidoreductases. For example, catalase is an important cellular antioxidant enzyme that protects against oxidative stress and catalyzes the decomposition of hydrogen peroxide to water and oxygen. Catalase is commonly present in nature (66), and here catalase had the greatest abundance in chestnut soil. Similarly, superoxide dismutase also protects against reactive oxygen species. Superoxide dismutases are metalloenzymes that catalyze the dismutation of iron superoxide into oxygen and hydrogen peroxide. The superoxide radical production increases under abiotic and biotic stresses, including drought (67) and had the lowest abundance in chestnut soil. Overall, differences in gene abundance related to stress response were slight.

### Potential limitations of the study

Shotgun metagenomic sequencing, as opposed to microarray and PCR analyses, is largely considered to yield unbiased results of gene presence and abundance (68, 69) because it does not require amplification prior to sequencing. However, the ability to interpret results for genes that are present in low abundance (here, *amoA, nifH, hzo, nirS, narB*) may become constrained (28, 55), although these genes may still provide function to microbial metabolism. This sequence-based approach may mask the importance of low abundance genes, as dominant populations may be oversampled, resulting in underestimation of low abundant genes (55). To improve interpretation for low abundance gene families, it is suggested that greater sequencing depth is necessary (28) and shotgun metagenomic sequencing should be used in conjunction with additional microarray or qPCR techniques to acquire both relative abundance of functional genes and capture the diversity of poorly represented families. Other potential limitations that may occur in metagenomic sampling may result from inadequate number of samples (spatially, temporally) that may not represent the full microbial community (70). We present results for one point in time and attempted to capture spatial variability by creating homogenized, composite samples from 10 points across each sampling plot.

## Conclusions

The results of our metagenomic analysis indicate that tree species can mediate the abundance of multiple key functional genes that encode for enzyme production in N metabolism pathways, and to a lesser degree for C metabolism, in soil. American chestnut exhibited the most distinct soil microbiome, both functionally and taxonomically, with a general suppression of N-cycling functional genes related to low inorganic N availability as soil was modified by poor, low N litter quality in chestnut relative to black cherry and northern red oak. This study provides one of the first comparative metagenomic analysis of the soil microbiome as it is modified by particular tree species. Our results provide insight into the mechanisms and degree of change in soil N and C metabolism that may occur following changes in dominant forest tree species through species loss by disease, management, or shifts due to a changing climate.

## Acknowledgements

This work was supported by a USDA NIFA McIntire-Stennis grant to CNK [award number 1008506], and a DOI Joint Genome Institute award to CNK and TPD [award number CSP 503303]. TPD was supported by start-up funding provided by West Virginia University. The funders had no role in study design, data collection and interpretation, or the decision to submit the work for publication.

The authors wish to thank Douglass Jacobs and Brian Beheler of Purdue University for access to tree plantations and field assistance, and Miranda Harmon-Smith for project management assistance with sample processing at JGI.

## Supplemental Material

All supplemental data for this study can be downloaded from the chestnut_metagenome repository on github (https://github.com/wvuvectors/soil_metagenome).

**Table S1.** Metadata for study samples present in the Integrated Microbial Genomes/Metagenomes database.

**Table S2.** Normalized abundance values for 36 genes involved in (A) nitrification, (B) denitrification, (C) nitrate reduction, and (D) annamox, N fixation, and N transport. All values were normalized to total metagenome CDS counts and multiplied by 10^6^.

**Table S3.** Normalized abundance values for 7 genes involved in organic N metabolism. All values were normalized to total metagenome CDS counts and multiplied by 10^6^.

**Table S4.** Normalized abundance values for 16 genes involved in C metabolism. All values were normalized to total metagenome CDS counts and multiplied by 10^6^.

**Table S5.** Normalized abundance values for 12 genes involved in stress response. All values were normalized to total metagenome CDS counts and multiplied by 10^6^.

**Table S6.** Gene names and database identifiers used in this study. EC: Enzyme Classification; KO: Kyoto Encyclopedia of Genes and Genomes. Colors signify functional group as defined in **Fig. 1**.

